# Burst firing and spatial coding in subicular principal cells

**DOI:** 10.1101/303354

**Authors:** Jean Simonnet, Michael Brecht

**Affiliations:** Bernstein Center for Computational Neuroscience Berlin, Humboldt-Universität zu Berlin, Philippstr. 13, Haus 6, 10115 Berlin, Germany; NeuroCure Cluster of Excellence, Humboldt-Universität zu Berlin, Berlin, Germany

**Author notes:** Corresponding Authors:Jean Simonnet, Michael Brecht, Bernstein Center for Computational Neuroscience Animal Physiology / Systems Neurobiology and Neural Computation Humboldt University Berlin Philippstr. 13, House 6 10115 Berlin, Germany.

**Keywords:** Hippocampus, orientation, cluster analysis, multiplexing

## Abstract

The subiculum is the major output structure of the hippocampal formation and is involved in learning and memory as well as in spatial navigation. Little is known about how the cellular diversity of subicular neurons is related to function. Primed by *in vitro* studies, which identified distinct bursting patterns in subicular cells, we asked how subicular burst firing is related to spatial coding *in vivo.* Using high-resolution juxtacellular recordings in freely moving rats, we analyzed the firing patterns of 51 subicular principal neurons and distinguished two populations based on their bursting behavior, i.e. sparsely bursting (∼80%) and dominantly bursting neurons (∼20%). Dominantly bursting neurons had significantly higher firing rates than sparsely bursting neurons. Furthermore, the two clusters had distinct spatial properties, sparsely bursting cells showing strong positional tuning and dominantly bursting cells being only weakly tuned. Additionally, the occurrence of bursts in sparsely bursting neurons defined well-defined spatial fields. In contrast, isolated spikes contained less spatial information. We conclude that burst firing distinguishes subicular principal cell types and constitutes a distinct unit encoding spatial information in sparsely bursting spatial cells. Overall, our results demonstrate that burst firing is highly relevant to subicular space coding.

## Significance statement

The subiculum is the major output structure of the hippocampal formation and is involved in spatial navigation. *In vitro,* subicular cells can be distinguished by their ability to initiate bursts, being brief sequences of spikes fired at high frequency. Little is known about the relationship between the cellular diversity and spatial coding in this structure. We performed high-resolution juxtacellular recordings in freely moving rats and found that bursting behavior predicts functional differences between subicular neurons. Specifically, sparsely bursting cells have lower firing rates and carry more spatial information than dominantly bursting cells. Additionally, bursts fired by sparsely bursting cells encoded spatial information better than isolated spikes, pointing towards bursts as a unit of information dedicated to space coding.

## Introduction

The subiculum is the major output structure of the hippocampus, receiving its main inputs from CA1 and sending divergent outputs to many subcortical and cortical areas (1, 2). The subiculum is involved in spatial learning and memory (3–5) but has not been the major focus of studies analyzing hippocampal function in spatial navigation.

*In vivo*, the vast majority of subicular neurons carry positional information - in various discharge patterns such as place fields, irregular spatial cells or boundary cells (6, 7). In addition, firing fields of subicular cells do not remap in response to novel environments, nor do they remap in darkness (6, 8, 9). A subset of subicular cells maps the current trajectory taken in an environment with well-defined routes rather than in an open-field arena, implying that environmental constraints strongly influence subicular cell coding (10). Subicular cell spatial fields are less well-defined than CA1 place cells (11), or the eye-catching medial entorhinal grid cells (12), due to higher basal firing rates in the subiculum compared to other spatial areas such as CA1 or medial entorhinal cortex.

The microcircuitry underlying spatial tuning in the subiculum is largely unresolved. The subicular anatomy is not as clearly stratified as the *stratum pyramidale* of CA1 (proximal to subiculum) and also lacks the elaborate lamination of the 6-layered cortical structures such as the presubiculum (distal to subiculum). The analysis of cell morphology indicates some internal structure (13) as well as laminar or modular organization based on long-range connectivity (2, 14–16). *In vitro,* subicular principal neurons may be distinguished by their firing patterns: some are intrinsic bursting (from 45 - 80%) and others are regular spiking cells (17, 18). Bursting relates to subicular anatomy: deeper cells as well as cells located on the distal part tend to be the most bursty (15, 17, 19). However, how bursting relates to subicular function remains mostly unresolved, even though a few functional correlates of bursting have been suggested. First, the biophysical properties of subicular cells could be predicted by their efferent target area (15), suggesting that intrinsic bursting or regular spiking cells might generate different streams of information. Second, local connectivity and recruitment by sharp wave ripples suggested distinct roles for regular spiking and intrinsic bursting cells in the subicular microcircuit function (20).

Here, we asked how subicular bursting relates to spatial coding *in vivo*. We took advantage of high-resolution juxtacellular recordings, which enabled us to reliably resolve small amplitude spikes - especially those resulting from sodium-channel inactivation during bursts. Using this technique in freely moving rats we asked: (i) Can bursting patterns be used to classify subicular neurons *in vivo* as *in vitro?* (ii) How does the burstiness of discharges relate to spatial coding? (iii) Do bursts and isolated spikes convey different types of information? We could classify cells based on their burstiness and found that sparsely bursting cells are more spatially modulated than dominantly bursting cells. In a large fraction of spatially modulated neurons, we found that bursts encoded position significantly better than isolated spikes. The encoding of spatial position by bursts predicts that such information is transmitted more effectively by facilitating synapses to downstream areas.

## Results

We performed juxtacellular recordings in rats foraging for food in a 70 × 70 cm open field arena. Our data consists of 51 subicular principal cells recorded in 28 rats. Neurons were assigned as subicular cells histologically and as principal cells based on their spike waveforms (See methods, Fig. S1).

### Sparsely and dominantly bursting cells: distinct firing patterns *in vivo*

Previous *in vitro* work (15, 17–19) indicated the existence of distinct patterns of bursting in different types of subicular principal cells. In our *in vivo* recordings, we also noted distinct bursting patterns of subicular principal cells during navigation. We categorized cells according to their burst discharge pattern. Clustering neurons using the Inter-Spike-Interval (ISI) histograms and spike autocorrelograms (Fig. S2; see methods) resulted in two distinct groups: sparsely bursting cells (n = 41 / 51 cells, ca. 80 %) and dominantly bursting cells (n = 10 / 51 cells, ca. 20 %; Fig. 1)

A trace from a recording and spikes of a sparsely bursting cell are shown in Fig. 1A and B. Burst firing (i.e. spikes interleaved with ISI ≤ 6 ms) occurred only sparsely. Both the ISI histogram (Fig. 1C) and autocorrelation function of spikes (Fig. 1D) confirm that short ISIs were present but rare in this cell group.

**Fig. 1.**
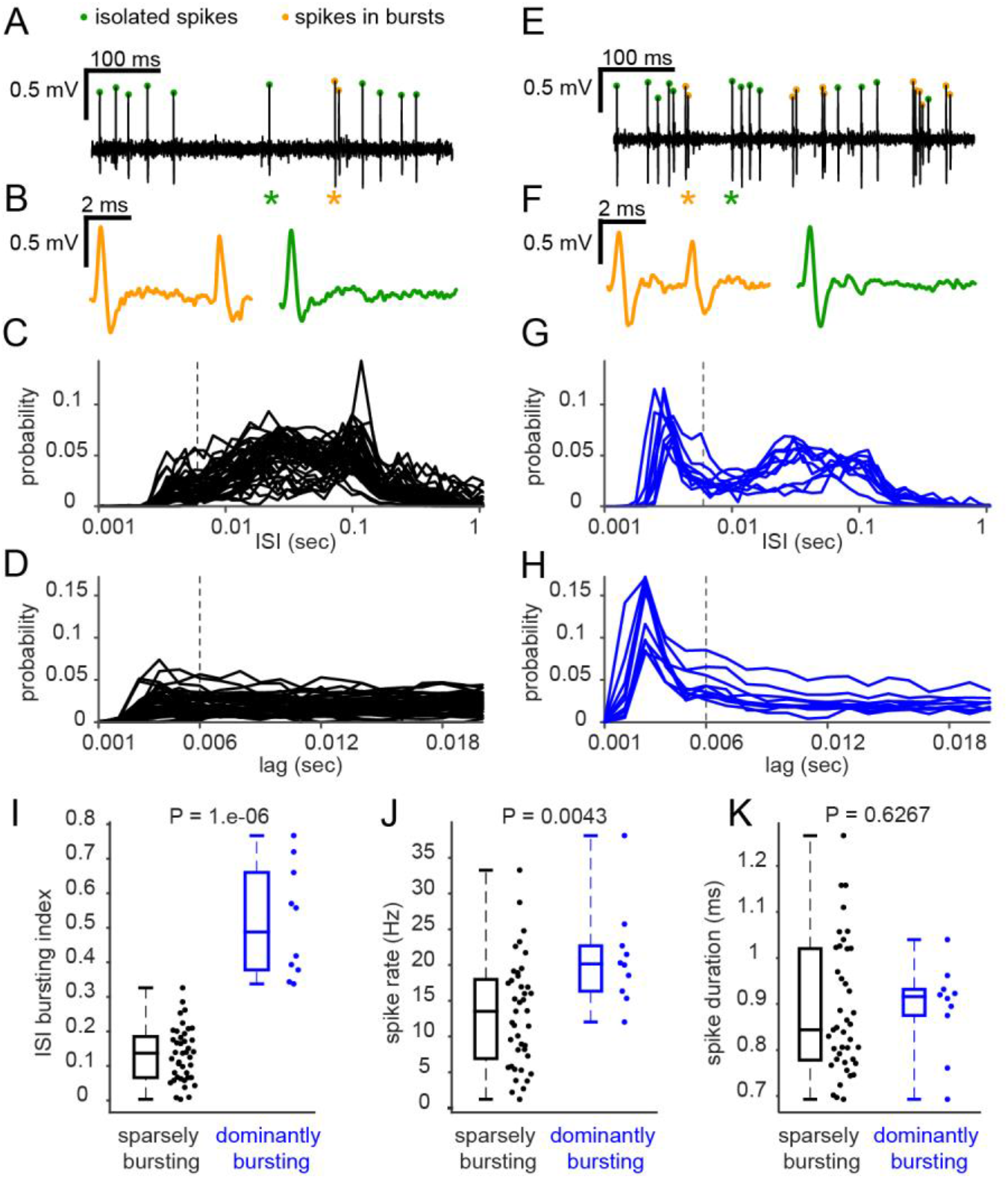
Firing pattern features of sparsely and dominantly bursting cells in vivo. (*A*) Bandpass filtered (300 - 6000Hz) trace of recording from a sparsely bursting cell. Spikes occurring in burst are labeled with an orange dots and isolated spikes with a green dots. (*B*) Magnification of the burst (orange, left) and the isolated spikes (green, right) indicated with a star. (*C*) Plot of the probability of Inter-Spike-Intervals (ISIs) for sparsely bursting subicular cells on a logarithmic scale. Note that short ISIs occur rarely in these neurons. The vertical dashed line is placed at 6 ms. (*D*) Autocorrelograms of sparsely bursting subicular cells. (*E, F*) as *B* and *C* for a dominantly bursting subicular cell. (*G*) Inter-Spike-Intervals as in C for dominantly bursting cells. Note the prominent initial peak. (*H*) Autocorrelograms of dominantly bursting subicular cells. (*I*) ISI based bursting index corresponding to the proportion of inter-spike-intervals lower than 6 ms is significantly higher for dominantly bursting cells. Two-tailed Mann Whitney U-Test. (*J*) Spiking rate (Hz) while the animal is navigating (see methods) is significantly higher for dominantly bursting cells. Two-tailed Mann Whitney U-Test. (*K*) Spike duration is not different between sparsely bursting and dominantly bursting cells. Statistics: two-tailed Mann Whitney U test.

A trace from a recording and spikes of a dominantly bursting cell are shown in Fig. 1E and F. A large fraction of spikes was organized in bursts. A prominent peak at short ISIs is obvious in the ISI histogram (Fig. 1G) and autocorrelation function of spikes (Fig. 1H).

The proportion of ISIs ≤ 6 ms of the total number of spikes was calculated as a bursting index. The median bursting index was significantly higher for dominantly bursting cells than for sparsely bursting cells (Fig. 1I, median: SB = 0.137, DB = 0.0.514, Mann-Whitney U test, P = 1.10^−6^). The average number of intra-burst spikes tended to be higher in dominantly bursting cells than in sparsely bursting cells (median: SB = 2.1, DB = 2.23, Mann-Whitney U test, P = 3.10^−4^). In addition, the mean intra-burst intervals were shorter in dominantly bursting cells than in sparsely bursting cells (median: SB = 4.3 ms, DB = 3.6 ms, Mann-Whitney U test, P = 2.10^−6^). Dominantly bursting cells had higher bursting rates than sparsely bursting cells (medians: SB = 0.7 Hz, DB = 4.8 Hz; Mann-Whitney U test, P = 3.10^−6^). Firing rates were variable and rather high as previously reported for subicular neurons (6, 7, 21). Firing rates of dominantly bursting cells were higher on average than sparsely bursting cells during navigating periods (speed > 1 cm.sec^−1^; Fig. 1J; medians: SB = 13.5 Hz, DB = 20.1 Hz; Mann-Whitney U test, P = 0.0043). Spike duration (from threshold to after-hyperpolarization) of sparsely bursting cells and dominantly bursting cells were not different from one another (Fig. 1K; median: SB = 0.85 ms, DB = 0.92 ms; Mann-Whitney U test, P = 0.6267).

### Sparsely bursting cells provide more spatial information than dominantly bursting cells

Our initial analysis showed that subicular cells can be clustered into two distinct cell populations based on their bursting activity *in vivo.* Next we asked if these populations also differ in their spatial tuning properties.

Fig. 2 shows the animals’ running trajectory with the superimposed spike positions and the resulting rate maps for one sparsely bursting cell (Fig. 2A, B) and one dominantly bursting cell (Fig. 2C, D). The sparsely bursting cell is robustly spatially tuned, while the dominantly bursting neurons is far less tuned. We calculated the spatial information for cells with sufficient coverage (>60%) of the open field area. This analysis (see methods, Fig. 2E) revealed that sparsely bursting cells are far more spatially tuned than dominantly bursting neurons (Fig. 2E, median: SB = 0.19 bits/spike (n = 35), DB = 0.03 bits/spike (n = 6), Mann-Whitney U test, P = 0.028). Given the size difference between groups and the low number of dominantly bursting cells (n = 6), we further tested the significance of the difference using bootstrapping (see methods). The spatial information of dominantly bursting cells was significantly different from the distribution of spatial information values of the randomly selected subsets of the sparsely bursting cells (P = 0.01).

**Fig. 2.**
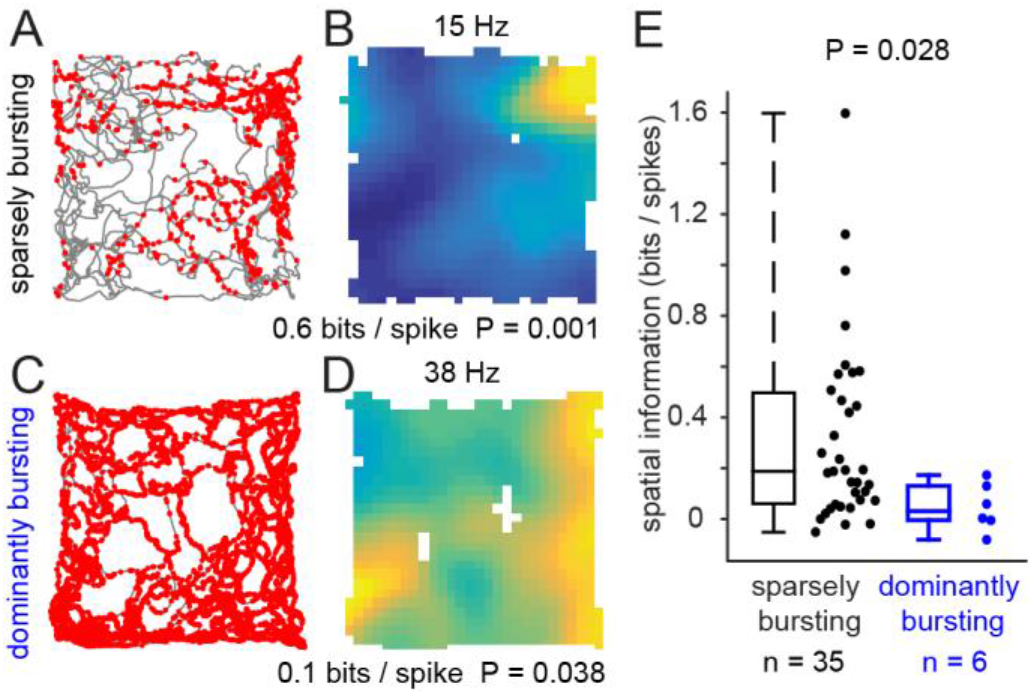
Sparsely bursting cells provide more spatial information than dominantly bursting cells. (*A*) Trajectories of the rat with the spikes of a sparsely bursting cell superimposed in red. (*B*) Corresponding rate map with the peak rate (above) and spatial information and significance (below). (*C*) As *A* for a dominantly bursting cell. (*D*) As *B* for a dominantly bursting cell. (*E*) Spatial information calculated for sparsely bursting and dominantly bursting cells. Sparsely bursting cells provided significantly more information than dominantly bursting cells. Statistics: two-tailed Mann Whitney U test.

Approximately 65% (25/35) of sparsely bursting cells and 50% (3/6) of dominantly bursting cells were significantly modulated by the animal’s position and could therefore be defined as spatial neurons. Sparsely bursting spatial neurons encoded more spatial information than dominantly bursting neurons (median: SB = 0.42 bits/spike, n = 25; DB = 0.13 bits/spikes, n = 3, bootstrapping P = 0.04).

The differences in spatial coding between sparsely bursting and dominantly bursting cells reinforced the idea that the classification of cells according to bursting discharge patterns captures significant functional differences between subicular neurons.

### Bursts provide spatial information in sparsely bursting cells

Our previous analyses revealed that subicular cells can be clustered according to bursting discharges and that such a classification reveals functional differences. While dominantly busting cells burst most of the time, sparsely bursting cells do so occasionally and we wondered, how such occasional bursts contributed to the transmission of spatial information. In the example shown in Fig. 2A and B, it can be seen that many spikes occurred outside of the main spatial firing field. In order to evaluate how the firing patterns contributed to the coding of spatial information at the single cell level, we separated isolated spikes and bursts into distinct plots (Fig 3A-F). A first example from a sparsely bursting cell, shown in Fig. 3A-C, suggested that isolated spikes occurred in numerous locations, even though a preferred location is still evident in the rate map (Fig 3B). A strikingly different picture emerged when we only plotted bursts. A well-defined firing field akin to a CA1 place cell emerged by looking at the bursts’ positions on the trajectory and the corresponding rate map (Fig. 3C). Applying the same analysis to further cells strengthened the idea that sparsely bursting cells provide spatial information through bursts. As shown for a second sparsely bursting cell in Fig. 3D-F, bursts were confined to the north border of the environment (Fig 3F) whereas isolated spikes were widely distributed throughout the arena (Fig 3E).

**Fig. 3:**
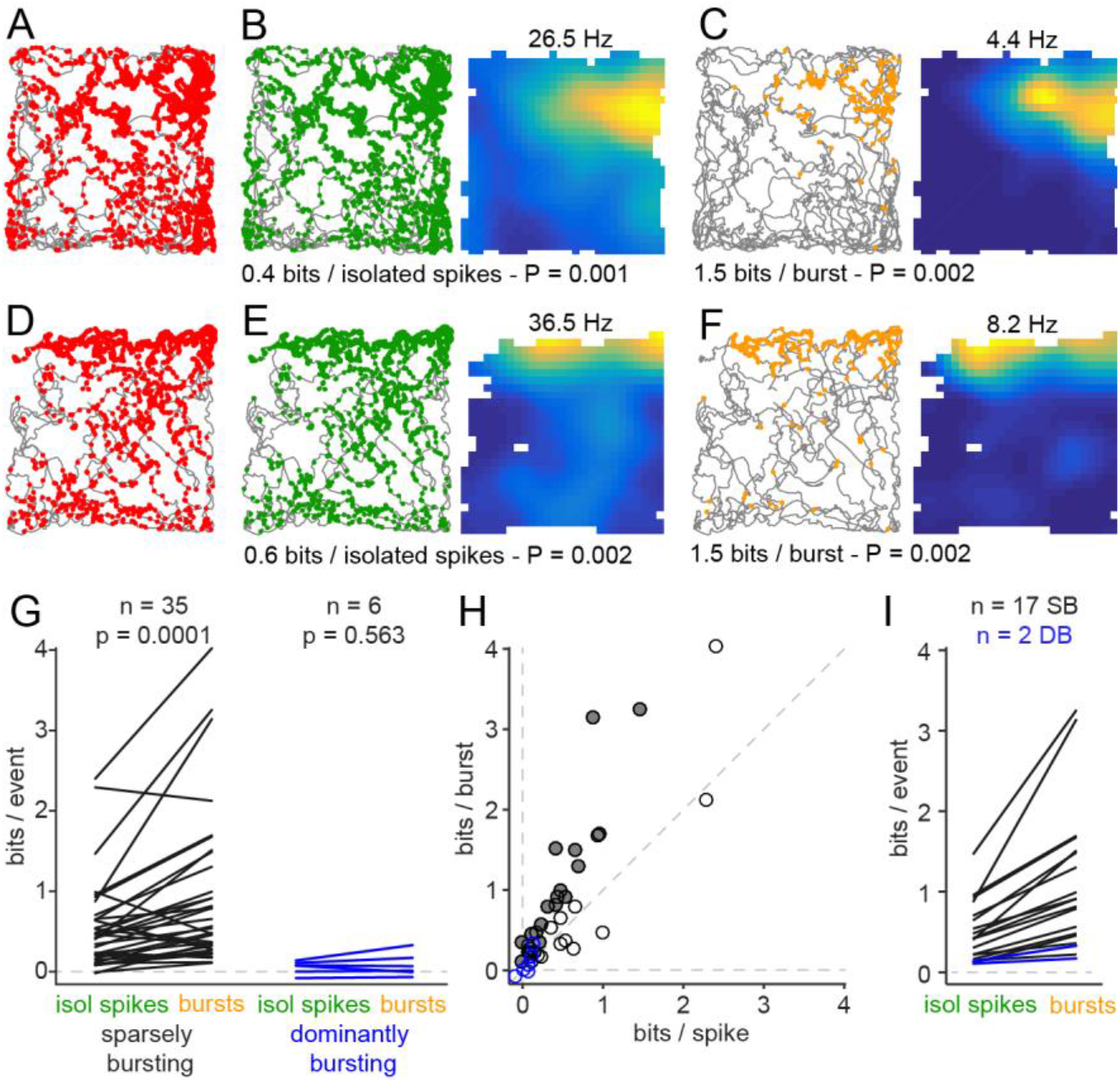
Sharp spatial tuning of burst firing in sparsely bursting cells. (*A*) Spikes (red dots) superimposed to the animal’s trajectory (gray line). (*B*) Left, isolated spikes (green dots) superimposed to the animal’s trajectory (gray line) and corresponding rate map; peak rate above and spatial information and significance below. (*C*) Left, bursts (orange dots) on animals’ trajectories (gray line) and corresponding burst rate maps; peak rate above and spatial information and significance below. (*D-F*), as *A-C* for another sparsely bursting cell. (*G*) Spatial information is significantly higher for bursts than isolated spike spatial information within the population of sparsely bursting cells but not for dominantly bursting cells. P, two-tailed Mann-Whitney U test. (*H*) Information per burst versus information per isolated spikes for spatially modulated neurons (n = 25 sparsely bursting cells, black circles; n = 3 dominantly bursting cells, blue circle). Solid circles indicate cells with a significant increase of spatial information between burst and isolated spike (n = 17 / 25 for sparsely bursting cells and n = 2 / 3 for dominantly bursting cells). (*I*) Same as *G*, showing only spatial cells with a significant difference between information encoded by bursts and isolated spikes, with sparsely bursting cells in black and dominantly bursting cells in blue. In *G-I,* spatial information values have been calculated using similar numbers of bursts and isolated spikes in order to obtain unbiased and comparable results (see methods).

The difference in the number of events per group (bursts or isolated spikes) can bias spatial information values; it tends to be higher while computed on a lower number of events (22). Typically, there was more isolated spikes than bursts. We compared the value of the spatial information encoded by the N bursts to the average of the spatial information encoded by 1000 random subsets of N isolated spikes (or the opposite if there was more isolated spikes than bursts). In sparsely bursting cells, burst transferred higher spatial information than isolated spikes (medians: 0.51 bits / burst, 0.41 bits / isolated spike; Wilcoxon signed rank test, P = 0. 0001; Fig. 3G). In contrast, bursts did not encode sharper spatial information in dominantly bursting cells (medians: 0.03 bits / burst, 0.07 bits / isolated spike; Wilcoxon signed rank test, P = 0.84; Fig. 3G). Here the median difference appears to be low. However, this analysis considered all neurons of our dataset, including non-spatial neurons. The median difference between bursts and isolated spikes was higher if we considered spatially significant sparsely bursting cells only (n = 23, 0.80 bits / burst, 0.44 bits / isolated spike; Wilcoxon signed rank test, P = 0.0001).

At the single cell level, information encoded by bursts was determined to be significantly more informative than isolated spikes if it was within the 95^th^ percentile of the distribution of spatial information values random subsets of N isolated spikes (See methods for when there are fewer isolated spikes than bursts). The difference between isolated spikes and bursts was significant for 17 of the 23 spatially modulated sparsely bursting cells and 2 of the 3 spatially modulated dominantly bursting cells (Fig. 3G, H). For sparsely bursting cells with a significant difference (n = 17), burst spatial information (median = 0.912 bits / burst) was on average 2.4 times higher than isolated spike spatial information (median, 0.42 bits / isolated spike, Fig. 3I). Similarly, it was 2.4 and 1.6 times higher for the two dominantly bursting spatial cells with a significant burst effect, nevertheless the burst spatial information remained among the lowest from our dataset (0.33 and 0.17 bits / burst, Fig. 3I).

## Discussion

We studied how burst firing related to spatial coding in the subiculum of rats. We first classified subicular neurons according to their bursting patterns and distinguished two classes of subicular neurons, a large fraction (80%) of sparsely bursting cells and a small fraction (20%) of dominantly bursting cells. Most sparsely bursting cells were spatially modulated cells and carried more spatial information than dominantly bursting cells. Finally, we found that bursts in sparsely bursting neurons carry more spatial information than isolated spikes, which account for the ubiquitous firing seen in most of the subicular spatial cells.

The initial impetus for our study came from *in vitro* studies, which identified bursting in subicular neurons. Most interestingly, bursting was shown to be correlated with the projection target of the respective neuron (15) suggesting a functional relevance of the bursting phenotype. In previous studies, bursting relationship to space coding has been investigated in subiculum without establishing a clear picture (6–8, 21). Previous reports on CA1 place cells suggested than intrinsic bursting cells, rather than regular spiking cells, were more likely to be spatially modulated (23). Unlike previous studies on the subiculum, we observed marked functional differences between cell classes defined by their bursting behavior, however quite opposite to CA1. Indeed, we found that dominantly bursting cells fire at higher rates and their spikes carry little spatial information, greatly strengthened the idea that bursting is of functional significance for subicular space coding. We believe that the high resolution of the juxtacellular recordings as well as our method to cluster subicular cells in distinct groups reflecting burstiness were key element in our findings.

However, it is not yet clear how our *in vivo* classification of sparsely bursting and dominantly bursting cells is related to various classifications of bursting based solely on intrinsic properties. Different *in vitro* studies on subicular neurons reported varying estimates for the fraction of intrinsically bursting neurons, ranging between 45 to 80 *%* (15, 17–19, 24–27). Only about 20% of the neurons observed in our study were of the dominantly bursting subtype. These numbers do not match previous reports and it seems unlikely that the dominantly bursting cells observed here correspond to the broad definition of intrinsic bursting cells used in *in vitro* studies. It seems possible that the dominantly bursting cells observed by us correspond to a subgroup of neurons with a particular strong tendency for intrinsic bursting described *in vitro.* In contrast, generating bursts does not appear to be a default mode of firing for most sparsely bursting cells. These cells might be weakly bursting or regular spiking cells requiring more complex mechanisms such as the interaction of intrinsic mechanisms and synaptic inputs for bursting (28).

Matching numbers of bursting cells between distinct studies is complicated because experimental conditions might be different from one study to another, especially since burst generation depends on many factors, which are not easy to control *in vivo.* Indeed, a variety of cellular mechanisms of burst generation have been suggested for subicular neurons. For instance, bursting requires T-type voltage gated calcium currents (25) that can be affected by neuromodulatory signals, such as serotonin, which was shown to downregulate T-type channels and burst generation (29).

As the output structure of the hippocampus, the subiculum sends high frequency, but rather unprecise, spatial coding to downstream areas. Indeed, peak frequencies of subicular spatial neurons are rather high compared to CA1, as are their baseline ((9, 15, 30). Nevertheless, spatial signals can be refined if one take the precise firing pattern of subicular neurons into consideration. Indeed, isolated spikes and bursts are functionally distinct units of information in most sparsely bursting spatial neurons (ca. 70%). While bursts were often fired in well-defined place fields, isolated spikes were spatially dispersed. Such differential coding by isolated spikes and bursts is similar to information processing in sensory systems (31). Nonetheless, our finding is remarkably different from CA1, where bursts sharpen spatial information in only ca. 20% of place cells (22). Such a difference shows the relevance of burst firing in noisy spatial cells such as subicular cells, compared to sharply tuned CA1 place cells. Bursts and isolated spikes, as two units of information could be readout by the interaction between short-term plasticity and postsynaptic integrative properties (32, 33). The spatial information conveyed by a burst could be decoded by the summation of excitatory events at facilitating synapses whereas poorly tuned spatial inputs could be better decoded through depressing synapses (33). This should be the case for long range projections and could as well define functional sub-circuits within the local microcircuit (34). The ongoing activity and resonating properties of targeted neurons could define the response to these signals (32). However, the neuronal targets of subicular spatial neurons and how these integrate and convert multiplexed signals at the cellular and microcircuit levels are unknown elements. These will need to be resolved for a better understanding of the subicular role in distributing hippocampal output spatial codes.

## Methods

All experimental procedures were performed according to German guidelines on animal welfare.

### Juxtacellular recordings in freely moving rats

Experimental procedures for obtaining juxtacellular recordings in freely moving rats were performed similar to earlier publications (35). Recordings were made in 28 male Long-Evans rats (150-350 g) maintained in a 12-h light / dark phase and recorded during the dark phase. Surgical procedures were all performed under ketamine (80-100 mg.kg^−1^) and xylazine (8010 mg.kg^−1^) anesthesia. Rats were implanted with a head-implant including a metal post for head-fixation, a placement of a miniaturized preamplifier coupled to two LEDs (red and blue) and a protection cap. In order to target the dorsal subiculum, a plastic ring was glued on the skull surface 5.7-6 mm posterior to bregma and 2.9-3.2 mm left to midline. The craniotomy and the positioning of the metal post for holding the miniaturized micromanipulator (Kleindiek Nanotechnik GmbH, Kusterdingen, Germany) were done either during the same surgery or in a subsequent surgery. After implantation, rats were allowed to recover and were habituated to head-fixation for 2-5 days. Rats were trained to forage for chocolate pellets in an open field arena - a 70 × 70 x 50 cm (WDH) box with a white polarizing cue card on one of the walls - prior to and after implantation (3-7 days, multiple sessions of 15-20 min each per day). For recordings, rats were head-fixed and the miniaturized micromanipulator and preamplifier were secured to the metal posts.

Glass pipettes with resistance 4-6 MQ were filled with Ringer solution (n = 45/53) containing (in mM) 135 NaCl, 5.4 KCl, 5 HEPES, 1.8 CaCl_2_, and 1 MgCl_2_; or patch clamping internal solution (n = 6/53) containing (in mM) 130 K-gluconate, 10 Na-gluconate, 10 HEPES, 10 phosphocreatine, 4 MgATP, 0.3 GTP, 4 NaCl. In both cases, pH was adjusted to 7.2, Neurobiotin (1-2%) was added to the solution and the osmolality was adjusted to 285-305 mmol/kg. The patch clamp solution was used to perform juxtacellular stimulations, of which the results are not used in the context of the current study. The firing rate and the firing pattern was not different between subicular cells recorded with the 2 different solutions; therefore the two subsets have been merged and considered as one group.

The glass recording pipette was advanced into the brain; a thick agarose solution (3.5-4% in Ringer) was applied into the recording chamber for sealing the craniotomy and stabilization. Animals were then released into the behavioral arena and juxtacellular recordings were established while animals were freely exploring the environment. The juxtacellular signals were acquired with an ELC-03XS amplifier (npi electronic GmbH, Tamm, Germany) and digitized with a Power 1401 data-acquisition interface coupled to Spike2-v7 (CED, Cambridge Electronic Design, Cambridge, UK) where signals were sampled at 50 kHz. The arena was filmed from above with a color camera so the position of red and blue LEDs could be tracked offline to determine animal’s location and head-direction. All signal processing and analyses were performed in Matlab (MathWorks, Natick, MA, USA).

### Anatomy

Juxtacellular labeling was performed at the end of the recording session according to standard procedures (36). A number of recordings were either lost before the labeling could be attempted, or the recorded neurons could not be clearly identified, but the location of all the cells included in the current study was positively assigned to the subiculum. Ten to thirty minutes after the labeling protocol, the animals were killed by prolonged isoflurane exposure and overdose of urethane, and perfused transcardially with 0.1 M PB followed by 4% paraformaldehyde solution. We used standard procedures for histological analysis of juxtacellularly-labeled neurons. Neurobiotin labeling was visualized with streptavidin conjugated to Alexa 488 (1:250 to 1:1000). Fluorescence images were acquired and position of tracks, filled neurons or recording sites were assigned to an area (subiculum, CA1 or presubiculum).

### Spike and bursts detection

For spike detection, the raw signals were filtered (0.3 - 6 KHz, zero phase band-pass Butterworth filter of order 8). Transients were then detected using a threshold of 2.5 times the root mean square (rms) of the signal. High amplitude artefacts, due to behaviors like grooming, could increase the rms value significantly and prevent the detection of the smallest transients; the values in a window of 2.5 ms around these artefacts were therefore clipped and replaced by zeros. A second step for separating spikes from noise consisted of calculating the principal components of the transients followed by manually clustering the events to spikes and noise. This cleaning step was first performed on filtered waveforms and subsequently on raw waveforms. Eventually, the accuracy of spike detection was visually checked, scrolling throughout the whole recording. The cleaning step was repeated until the detection was optimal (minimizing false positives and negatives).

Finally, spikes were categorized as belonging to a burst if the interval from the prior spike and/or to the next spike was shorter than a threshold set at 6 ms. One burst was therefore defined as a group of spikes (>=2) interleaved with less than 6 ms. The burst time stamp was set to that of the first spike in a burst. Burst length was the time difference between the last and the first spike in a burst and burst modal interval was the mean inter-spike-interval (ISI) during bursts.

### Spike waveform analysis

The raw signals were filtered (6 KHz, zero phase low-pass Butterworth filter of order 8) in order to minimize high frequency noise. Spike shape parameters were determined based on the spike average waveform calculated from these low-pass filtered traces. Prior to the calculation of the average spike, the single waveforms had to be properly aligned. T o this end, every spike waveform was oversampled at 1000 kHz using a spline interpolation to better estimate its shape. We then calculated the derivative of each waveform and derived the threshold point as the first point where the derivative waveform first reached 5% of its maximum (i.e corresponding to depolarization rate reaching 5% of the maximal depolarization rate).

The rising amplitude (mV) was set to the difference of potential between the peak and the threshold voltage. Signal to noise ratios often differed between recordings and with it, the spike amplitude. To be able to compare spike shape parameters between cells the waveform was normalized so that the rising amplitude was 1 mV. The after-hyperpolarization was set to the point at minimum voltage after the peak. The spike duration was set to the threshold-to-after-hyperpolarization duration. Putative fast-spiking interneurons were identified based on their spike duration inferior to 0.5 ms (n = 2, 0.39 and 0.43 ms, Fig. S1) and not used for the subsequent analyses.

### Analysis of burstiness

We estimated the burstiness of each subicular principal cell using a combination of two different methods, which are both biased by basal firing rates but in opposite directions (Fig. S2).

The first method is based on the distribution of inter-spike-intervals and has previously been used in order to study a cell’s burstiness in the subiculum (6–8) and other areas such as parahippocampal cortices (37). The ISI interval distribution might overestimate burstiness for cells with elevated firing rates. This problem is solved by the second method - analysis of the spike autocorrelograms from 1 to 20 ms (21). Kim et al. (2012) have calculated a bursting index as the ratio of the integrated power of the autocorrelogram between 1 and 6 ms normalized by the overall power between 1 and 20 ms. This method does not bias the burstiness estimation for high firing neurons. However, we realized that it could overestimate the bursting probability of neurons with low firing rate and very occasional bursts because only the first 20 ms of the autocorrelogram are considered in this analysis.

Principal component analysis (PCA) was done on both the log(ISI) probability matrix and for the 1-20 ms lag probability matrix. The first three components from each PCA were used to generate a firing pattern vector in a 6-dimensional space. We then generated a cluster tree using Ward’s method on the normalized Euclidean distance between cells. The Ward’s method establishes hierarchical clusters by iteratively grouping the two closest observations or groups of observations together. Consequently, cells with very similar firing patterns are primarily grouped together and groups with very different properties are linked at the end of the procedure (38). Two clusters strikingly emerged from the dendrogram, defining two groups of neurons that we named sparsely bursting cells and dominantly bursting cells based on their potency to initiate bursts.

### Analysis of spatial modulation

The position of the rat was defined as the midpoint between two head-mounted LEDs. A running speed threshold (1 cm.sec^−1^) was applied for isolating periods of rest from navigation. For generating color-coded firing maps, space was discretized into pixels of 2.5 cm x 2.5 cm. For each such pixel the occupancy *o(x)* was calculated:

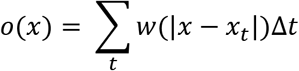

where *x_t_* is the position of the rat at time *t*, *Δt* the inter-frame interval, and *w* a Gaussian smoothing kernel with *σ* = 5 cm. Then, the firing rate *r* was calculated for each pixel x:

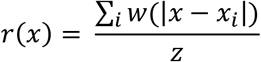

where *x_i_* is the position of the rat when spike *i* was fired.

For recordings in which the animal’s trajectory covered at least 60 % of the open field (n = 41/51), we calculated the spatial information rate, I (bits / spike), from the spike train and rat trajectory as follows:

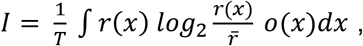

where r(x) and o(x) are the firing rate and occupancy as a function of a given pixel x in the rate map. 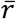 is the overall mean firing rate of the cell, and T is the total duration of a recording session (39).

## Statistics

A cell was declared to have a significant amount of spatial information if the observed spatial information rate exceeded the 95^th^ percentile of a distribution of values of I obtained by circular shuffling. Shuffling was performed a 1000 times by a circular time-shift of the recorded spike train relative to the rat’s trajectory by a random time *t’* ϵ [0 *T*] (40, 41), T being the total duration of the recording session.

The size difference between the two clusters of subicular neurons and the small size of the group of dominantly bursting cells used for spatial information calculation brought us to test the significance of the difference with a bootstrapping procedure. N (number dominantly bursting cells) values of spatial information were randomly selected from the sparsely bursting population. Repeating the procedure 1000 times, we then obtained a bootstrapped distribution of the median spatial information for sparsely bursting cells. The difference was significant if the rank of the median spatial information of dominantly bursting cells was within the 5^th^ percentile of the bootstrapped distribution (P ≤ 0.05).

We wanted to test whether the spatial information encoded by bursts was significantly different than the spatial information encoded by isolated spikes. A direct comparison of spatial information values would not be appropriate as the total number of events and smoothing parameters used for generating the rate maps can introduce bias in information-theoretic measures (22). Consequently, we used a randomization method similar to Harris et al. (2001b). In instances where there were less bursts than isolated spikes, we would compare information given by the N bursts to the information given by 1000 random subsets of N isolated spikes. Bursts were significantly more informative than isolated spikes if the rank of the burst spatial information was within the 95^th^ percentile of the distribution of the spatial information given by the random subsets of isolated spikes.

In some instances where highly bursting cells had less isolated spikes than bursts (n = 2/51), we compared the information given by N isolated spikes to the information given by 1000 random subsets of N bursts. In this case, bursts were significantly more informative than isolated spikes if the rank of the isolated spike spatial information was within the 5^th^ percentile of the distribution of the spatial information given by the random subsets of bursts.

In many cases, the significance of the difference observed between distinct groups was assessed with non-parametric tests only. Two-tailed Mann-Whitney U tests were used to determine whether two groups of unpaired observations were significantly different from each other (e. g. comparing sparsely bursting and dominantly bursting cells). Two tailed Wilcoxon signed rank tests were used in case of two groups of paired observations (e.g. comparing bursts and isolated spikes)

## Acknowledgements

This work was supported by Humboldt-Universität zu Berlin, BCCN Berlin (German Federal Ministry of Education and Research BMBF, Förderkennzeichen 01GQ1001A), NeuroCure, Neuro-Behavior ERC Grant, Deutsche Forschungsgemeinschaft Gottfried Wilhelm Leibniz Prize. We thank Edith Chorev, Rajnish Rao, Juan Ignacio Sanguinetti-Scheck and Peter Bennett for discussion and comments on the manuscript. We thank Andreea Neurkirchner, Undine Schneeweiß, Juliane Diederichs, Tanja Wölk, Maik Kunert and Arnold Stern for outstanding technical support.

## References

1. Amaral DG, Witter MP (1989) The three-dimensional organization of the hippocampal formation: a review of anatomical data. Neuroscience 31(3):571–591.

2. Witter MP (2006) Connections of the subiculum of the rat: Topography in relation to columnar and laminar organization. Behav Brain Res 174(2):251–264.

3. Galani R, Weiss I, Cassel J-C, Kelche C (1998) Spatial memory, habituation, and reactions to spatial and nonspatial changes in rats with selective lesions of the hippocampus, the entorhinal cortex or the subiculum. Behav Brain Res 96(1):1–12.

4. Morris RGM, Schenk F, Tweedie F, Jarrard LE (1990) Ibotenate lesions of hippocampus and/or subiculum: dissociating components of allocentric spatial learning. Eur J Neurosci 2(12):1016–1028.

5. Roy DS, et al. (2017) Distinct Neural Circuits for the Formation and Retrieval of Episodic Memories. Cell 170(5):1000–1012.e19.

6. Lever C, Burton S, Jeewajee A, O’Keefe J, Burgess N (2009) Boundary Vector Cells in the Subiculum of the Hippocampal Formation. J Neurosci 29(31):9771–9777.

7. Sharp PE, Green C (1994) Spatial correlates of firing patterns of single cells in the subiculum of the freely moving rat. J Neurosci 14(4):2339–2356.

8. Brotons-Mas JR, Montejo N, O’Mara SM, Sanchez-Vives MV (2010) Stability of subicular place fields across multiple light and dark transitions: Stability of subicular place fields. Eur J Neurosci 32(4):648–658.

9. Sharp PE (1997) Subicular cells generate similar spatial firing patterns in two geometrically and visually distinctive environments: comparison with hippocampal place cells. Behav Brain Res 85(1):71–92.

10. Olson JM, Tongprasearth K, Nitz DA (2017) Subiculum neurons map the current axis of travel. Nat Neurosci 20(2):170–172.

11. O’Keefe J, Dostrovsky J (1971) The hippocampus as a spatial map. Preliminary evidence from unit activity in the freely-moving rat. Brain Res 34(1):171–175.

12. Hafting T, Fyhn M, Molden S, Moser M-B, Moser EI (2005) Microstructure of a spatial map in the entorhinal cortex. Nature 436(7052):801–806.

13. O’Mara S (2005) The subiculum: what it does, what it might do, and what neuroanatomy has yet to tell us. J Anat 207(3):271–282.

14. Ishizuka N (2001) Laminar organization of the pyramidal cell layer of the subiculum in the rat. J Comp Neurol 435(1):89–110.

15. Kim Y, Spruston N (2012) Target-specific output patterns are predicted by the distribution of regular-spiking and bursting pyramidal neurons in the subiculum. Hippocampus 22(4):693–706.

16. Naber PA, Witter MP (1998) Subicular efferents are organized mostly as parallel projections: A double-labeling, retrograde-tracing study in the rat. J Comp Neurol 393(3):284–297.

17. Greene JR, Totterdell S (1997) Morphology and distribution of electrophysiologically defined classes of pyramidal and nonpyramidal neurons in rat ventral subiculum in vitro. J Comp Neurol 380(3):395–408.

18. Staff NP, Jung H-Y, Thiagarajan T, Yao M, Spruston N (2000) Resting and active properties of pyramidal neurons in subiculum and CA1 of rat hippocampus. J Neurophysiol 84(5):2398–2408.

19. Harris E, Witter MP, Weinstein G, Stewart M (2001) Intrinsic connectivity of the rat subiculum: I. Dendritic morphology and patterns of axonal arborization by pyramidal neurons. J Comp Neurol 435(4):490–505.

20. Bohm C, et al. (2015) Functional Diversity of Subicular Principal Cells during Hippocampal Ripples. J Neurosci 35(40):13608–13618.

21. Kim SM, Ganguli S, Frank LM (2012) Spatial Information Outflow from the Hippocampal Circuit: Distributed Spatial Coding and Phase Precession in the Subiculum. J Neurosci 32(34):11539–11558.

22. Harris KD, Hirase H, Leinekugel X, Henze DA, Buzsáki G (2001) Temporal interaction between single spikes and complex spike bursts in hippocampal pyramidal cells. Neuron 32(1):141–149.

23. Epsztein J, Brecht M, Lee AK (2011) Intracellular Determinants of Hippocampal CA1 Place and Silent Cell Activity in a Novel Environment. Neuron 70(1):109–120.

24. Behr J, Empson RM, Schmitz D, Gloveli T, Heinemann U (1996) Electrophysiological properties of rat subicular neurons in vitro. Neurosci Lett 220(1):41–44.

25. Joksimovic SM, et al. (2017) The role of T-type calcium channels in the subiculum: to burst or not to burst?: T-channels regulate neuronal excitability in the subiculum. J Physiol 595(19):6327–6348.

26. Mason A (1993) Electrophysiology and burst-firing of rat subicular pyramidal neurons in vitro: a comparison with area CA1. Brain Res 600(1):174–178.

27. Stewart M, Wong RK (1993) Intrinsic properties and evoked responses of guinea pig subicular neurons in vitro. J Neurophysiol 70(1):232–245.

28. Larkum M (2013) A cellular mechanism for cortical associations: an organizing principle for the cerebral cortex. Trends Neurosci 36(3):141–151.

29. Petersen AV, Jensen CS, Crépel V, Falkerslev M, Perrier J-F (2017) Serotonin Regulates the Firing of Principal Cells of the Subiculum by Inhibiting a T-type Ca2+ Current. Front Cell Neurosci 11:60.

30. Sharp PE (2006) Subicular place cells generate the same “map” for different environments: Comparison with hippocampal cells. Behav Brain Res 174(2):206–214.

31. Krahe R, Gabbiani F (2004) Burst firing in sensory systems. Nat Rev Neurosci 5(1):13–23.

32. Izhikevich EM, Desai NS, Walcott EC, Hoppensteadt FC (2003) Bursts as a unit of neural information: selective communication via resonance. Trends Neurosci 26(3):161–167.

33. Lisman JE (1997) Bursts as a unit of neural information: making unreliable synapses reliable. Trends Neurosci 20(1):38–43.

34. Simonnet J, et al. (2017) Activity dependent feedback inhibition may maintain head direction signals in mouse presubiculum. Nat Commun 8:16032.

35. Tang Q, Brecht M, Burgalossi A (2014) Juxtacellular recording and morphological identification of single neurons in freely moving rats. Nat Protoc 9(10):2369–2381.

36. Pinault D (1996) A novel single-cell staining procedure performed in vivo under electrophysiological control: morpho-functional features of juxtacellularly labeled thalamic cells and other central neurons with biocytin or Neurobiotin. J Neurosci Methods 65(2):113–136.

37. Ebbesen CL, et al. (2016) Cell Type-Specific Differences in Spike Timing and Spike Shape in the Rat Parasubiculum and Superficial Medial Entorhinal Cortex. Cell Rep 16(4):1005–1015.

38. Ward JH (1963) Hierarchical Grouping to Optimize an Objective Function. J Am Stat Assoc 58(301):236–244.

39. Skaggs WE, McNaughton BL, Gothard KM (1993) An information-theoretic approach to deciphering the hippocampal code. Advances in Neural Information Processing Systems, pp 1030–1037.

40. Bjerknes TL, Moser EI, Moser M-B (2014) Representation of Geometric Borders in the Developing Rat. Neuron 82(1):71–78.

41. von Heimendahl M, Rao RP, Brecht M (2012) Weak and Nondiscriminative Responses to Conspecifics in the Rat Hippocampus. J Neurosci 32(6):2129–2141.

